# Low-coverage sequencing cost-effectively detects known and novel variation in underrepresented populations

**DOI:** 10.1101/2020.04.27.064832

**Authors:** Alicia R. Martin, Elizabeth G. Atkinson, Sinéad B. Chapman, Anne Stevenson, Rocky E. Stroud, Tamrat Abebe, Dickens Akena, Melkam Alemayehu, Fred K. Ashaba, Lukoye Atwoli, Tera Bowers, Lori B. Chibnik, Mark J. Daly, Timothy DeSmet, Sheila Dodge, Abebaw Fekadu, Steven Ferriera, Bizu Gelaye, Stella Gichuru, Wilfred E. Injera, Roxanne James, Symon M. Kariuki, Gabriel Kigen, Karestan C. Koenen, Edith Kwobah, Joseph Kyebuzibwa, Lerato Majara, Henry Musinguzi, Rehema M. Mwema, Benjamin M. Neale, Carter P. Newman, Charles R. J. C. Newton, Joseph K. Pickrell, Raj Ramesar, Welelta Shiferaw, Dan J. Stein, Solomon Teferra, Celia van der Merwe, Zukiswa Zingela, NeuroGAP-Psychosis Consortium

## Abstract

**Background:** Genetic studies of biomedical phenotypes in underrepresented populations identify disproportionate numbers of novel associations. However, current genomics infrastructure--including most genotyping arrays and sequenced reference panels--best serves populations of European descent. A critical step for facilitating genetic studies in underrepresented populations is to ensure that genetic technologies accurately capture variation in all populations. Here, we quantify the accuracy of low-coverage sequencing in diverse African populations.

**Results:** We sequenced the whole genomes of 91 individuals to high-coverage (≥20X) from the Neuropsychiatric Genetics of African Population-Psychosis (NeuroGAP-Psychosis) study, in which participants were recruited from Ethiopia, Kenya, South Africa, and Uganda. We empirically tested two data generation strategies, GWAS arrays versus low-coverage sequencing, by calculating the concordance of imputed variants from these technologies with those from deep whole genome sequencing data. We show that low-coverage sequencing at a depth of ≥4X captures variants of all frequencies more accurately than all commonly used GWAS arrays investigated and at a comparable cost. Lower depths of sequencing (0.5-1X) performed comparable to commonly used low-density GWAS arrays. Low-coverage sequencing is also sensitive to novel variation, with 4X sequencing detecting 45% of singletons and 95% of common variants identified in high-coverage African whole genomes.

**Conclusion:** These results indicate that low-coverage sequencing approaches surmount the problems induced by the ascertainment of common genotyping arrays, including those that capture variation most common in Europeans and Africans. Low-coverage sequencing effectively identifies novel variation (particularly in underrepresented populations), and presents opportunities to enhance variant discovery at a similar cost to traditional approaches.

## Background

Over the last decade, genome-wide association studies (GWAS) have grown rapidly, deepening biological insights into a breadth of human diseases. Data for these studies are usually generated using GWAS arrays due to their cost effectiveness and the availability of commonly used analytical pipelines. These arrays typically genotype a fixed set of hundreds of thousands to millions of common variants genome-wide, and additional linked variants are then imputed using haplotype reference panels (Marchini and Howie, 2010). The utility of this approach varies across populations, however, because most GWAS arrays consist of variants that are most common in European ancestry populations. Even if arrays ascertained variants with similar frequencies across populations, reference data for imputation are also vastly Eurocentric (Huang et al., 2009; McCarthy et al., 2016; Wojcik et al., 2018).

Recognition of these biases in genomic infrastructure has driven concerted efforts to develop specialized, scalable arrays designed to capture variation common to different continental ancestries. For example, the Population Architecture using Genomics and Epidemiology (PAGE) Consortium designed the Illumina Multi-ethnic Genotyping Array (MEGA), a dense array of ~1.7 million variants, which aimed to improve performance for imputation across globally diverse populations (Wojcik et al., 2018). A significant portion of the ~660,000 variants on the Global Screening Array (GSA)--designed to decrease costs, increase scalability, and improve imputation accuracy in European populations--consists of a subset of variants from MEGA. Additionally, the Human Heredity and Health in Africa (H3Africa) Consortium developed a dense array of ~2.5 million variants specialized for the higher genetic diversity and smaller haplotype blocks in African genomes (Mulder et al., 2018). While these arrays all have potential benefits, an inherent weakness to their ascertained nature is that they cannot capture novel variants.

As sequencing costs have dropped, low-pass sequencing has been proposed as a similarly priced and unbiased alternative to GWAS arrays, for example in population genetics and polygenic score analysis (Alex Buerkle and Gompert, 2013; Gilly et al., 2018; Homburger et al., 2019; Pasaniuc et al., 2012; Pickrell, 2017). Sequencing offers several advantages: 1) variants are unascertained, meaning that the quality of data generated is inherently unbiased towards any particular population, 2) novel, population-specific variants can be detected and used to further advance the generation of haplotype reference panels, 3) DNA strand is unambiguous given the alignment of sequencing reads to a reference genome, and 4) non-human microbiome DNA can be captured and variation analyzed with certain DNA sampling procedures. These advantages are expected to be especially beneficial in non-European populations because corresponding reference data that support arrays are often lacking.

Here, we have generated high coverage whole genome sequencing data from populations vastly underrepresented in genetics research to compare data quality that would be produced by sequencing at various depths versus genotyping with several commonly used arrays. We have also compared the costs and analytical approaches that are feasible from each data generation approach. To compare data generation strategies, we included whole genomes that were sequenced as part of the Neuropsychiatric Genetics of African Populations Psychosis (NeuroGAP-Psychosis) study spanning five sites across four countries in eastern and southern Africa (Stevenson et al., 2019). These populations are of particular interest because humans originated in Africa, resulting in high levels of genetic variation and rapid linkage disequilibrium decay, highlighting the disproportionate informativeness of African genomes for human evolutionary studies and in pinpointing causal variants. Thus, accurately capturing genetic variation in these populations in an unbiased manner is particularly important for associating, resolving, and interpreting genetic associations while ensuring equitable translation of genetic technologies. Our results highlight that low-coverage sequencing can be a more appropriate data generation strategy than GWAS arrays for assaying genetic variation across globally diverse populations.

## Results

To compare genetic data quality from variable depths of sequencing versus commonly used GWAS arrays, we sequenced the whole genomes of participants from the NeuroGAP-Psychosis study to high coverage (target coverage of ≥30X per individual, mean coverage = 38X, all ≥ 20X, **Supplementary Figure 1**). This study consists of data from five geographical sites (N=91, with N ≥ 17 individuals per site) across eastern and southern Africa (**Table 1, Figure 1A-B**). Participants in these studies were chosen from a larger set of genotyped individuals based on ancestry patterns representative of the enrollment site. They come from a range of ethnic groups, with more than five individuals per NeuroGAP-Psychosis recruitment site reporting the following primary ethnicities: Amhara and Oromo from Addis Ababa, Ethiopia; Xhosa from Cape Town, South Africa; Mijikenda from Kilifi, Kenya; and the Kalenjin from Eldoret, Kenya (**Supplementary Table 1**). There was no predominantly reported primary ethnicity among the 18 individuals from Kampala, Uganda; rather, 11 different ethnic groups were reported among these individuals.

**Figure 1 –.**
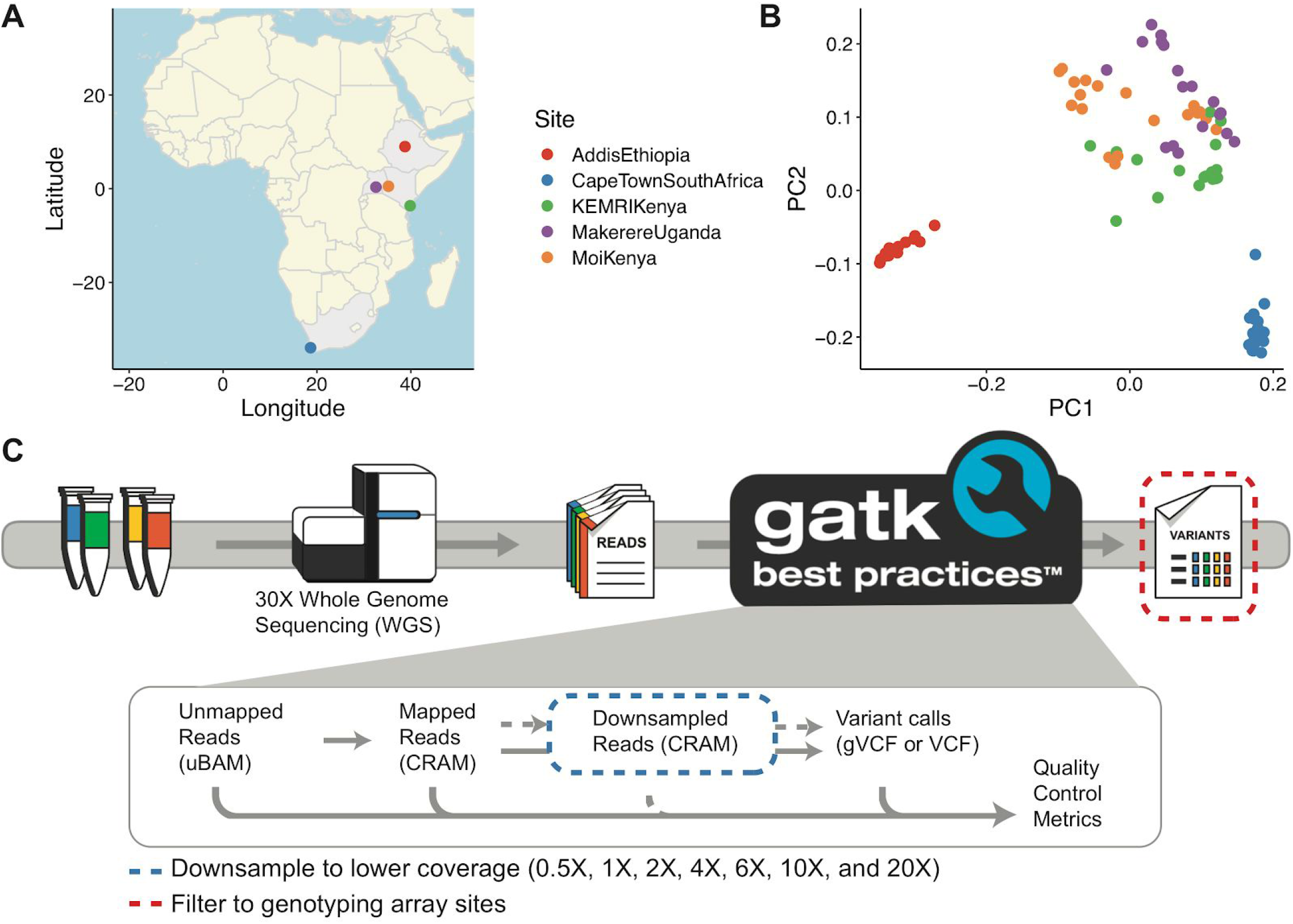
Populations and sites included in high coverage whole genome sequence data and downsampling schema to assess the performance of lower coverage sequencing versus GWAS arrays. A) Map indicating where participants in the NeuroGAP-Psychosis study are recruited from in this dataset. B) The first two principal components (PC) show variation within and among populations. They first distinguish the Ethiopians, then the South Africans from other African populations. Colors are consistent in panels A and B. C) High coverage genomes were processed using the GATK best practices pipeline. To mimic lower coverage sequencing data, analysis-ready CRAM files were downsampled to various depths, followed by a standard implementation of the variant calling pipeline. To mimic GWAS array data, the variants called from the high coverage sequencing data were filtered to only those sites on the arrays.

**Table 1.**
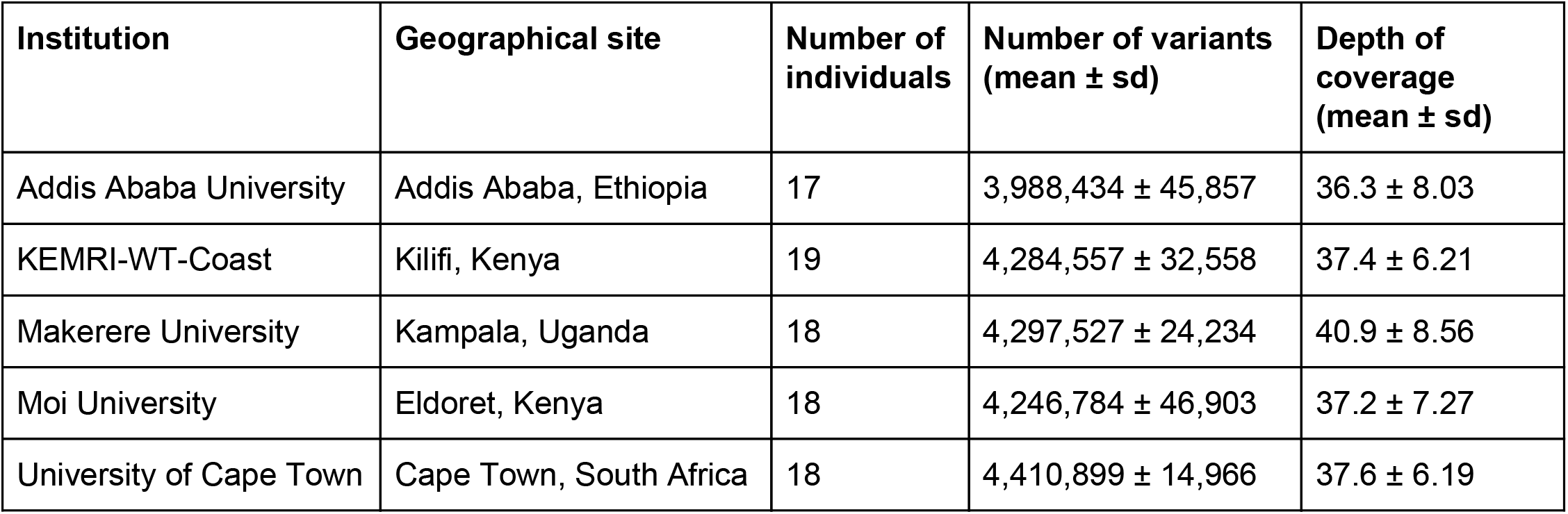
Genetic samples included in these sequencing analyses. 19 samples from Ethiopia were sequenced, but two showed significant evidence of contamination and were excluded from variant calling metrics and all downstream analyses. The number of variants reported are per-individual non-reference variant calls.

We considered variant calls generated from all reads to be our “truth” variant calls throughout our analyses. Across all individuals and geographical sites, these high coverage whole genomes contain 26 million variants, with more than 4 million non-reference variants per individual in all populations except in Ethiopia (**Table 1**). Consistent with our results, prior studies of Ethiopian genetics have shown reductions in genetic diversity compared to other African populations due to back-to-Africa migrations from the Middle East (Henn et al., 2012; Hodgson et al., 2014; Pagani et al., 2015).

We next downsampled or subset our data to simulate low-coverage and GWAS array data generation, respectively, using two approaches (**Figure 1C**). First, we downsampled analysis-ready CRAM files to the number of reads corresponding to 0.5X, 1X, 2X, 4X, 6X, 10X, and 20X coverage (**Methods**). With these downsampled data, we then generated new variant call sets corresponding to these depths (**Supplementary Table 2)** and performed variant QC using standard analysis pipelines (**Methods**). Second, we subset variants from the high coverage “truth” data corresponding to all polymorphic sites that would have been probed using each of the following Illumina arrays: the GSA, PsychChip, MEGA, H3Africa, and Omni2.5. For both of these datasets, we then compared the imputed data to the high coverage variant calls to assess the number and quality of sites obtained.

We first compared the downsampled versus highest depth “truth” whole genome sequencing data prior to any genotype refinement or imputation. Compared to high-coverage sequencing data, we expect low-coverage sequencing to produce variant calls that have higher error rates and miss some genetic variants altogether because of the reduced chance of observing both alleles with high quality reads across regions of the genome. We therefore calculated non-reference concordance (**Methods**) between the downsampled variant call sets and the full coverage data (**Figure 2, Supplementary Table 3**). Non-reference concordance was lower for indels than SNPs and was lowest for variants with ~5% frequency.

**Figure 2 –.**
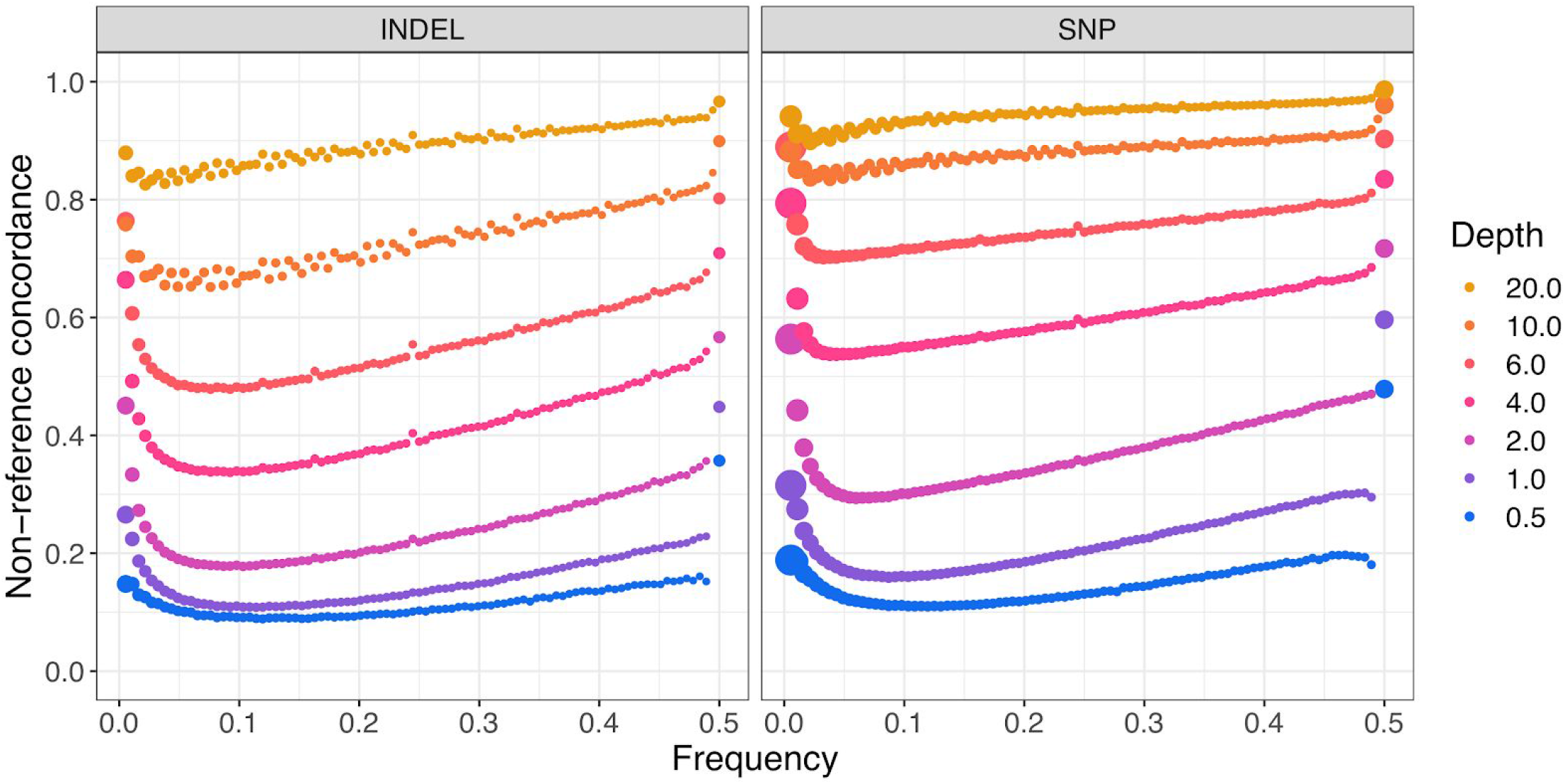
Pre-imputation non-reference variant concordance. We computed non-reference concordance comparing downsampled data at several depths of coverage to the highest depth sequencing call set available for all samples. The size of each dot is proportional to the number of variants in each bin. Depth summaries across samples are shown in **Supplementary Figure 1**. Non-reference concordances averaged across variants of all allele frequencies are shown in **Supplementary Table 3**.

We next used BEAGLE, open-source software described previously (Browning and Browning, 2007; Browning et al., 2018), for genotype refinement and imputation of low-coverage data (**Methods**)(CONVERGE consortium, 2015; Homburger et al., 2019; Luo et al., 2017). Genotype refinement is designed to correct low-quality genotype calls using a haplotype reference panel of high-confidence genotypes and considers genotype likelihoods rather than hard calls. Afterwards, imputation uses the refined genotype calls to fill in variants from the reference panel for sites not originally called. We performed genotype refinement and imputation on low-coverage sequencing up to 6X using 1000 Genomes phase 3 data as a haplotype reference panel (1000 Genomes Project Consortium et al., 2015). The higher depths, 10X and 20X, were excluded given their already high concordance without refinement (**Figure 2**) and to save computational costs. To compare variant calls obtained from our whole genome sequencing experiment with several commonly used genotyping arrays, we filtered variants from the high coverage “truth” dataset to those on the array, then imputed genotypes using the same methodology as in the downsampled sequencing data (**Figure 1B**).

We first compared non-reference concordance in the low-versus high-coverage sequencing data using variant calls through each step of the process, including the raw data, after genotype refinement, and after imputation (**Methods**). The total numbers of SNPs through each processing step are shown in **Table 2** (imputed > raw > refined). Prior to imputation, we identify approximately 13 million variants from 1X sequencing compared to the 26 million in the high coverage data (~50%). This is a considerably larger number of polymorphic variants than are genotyped on any array (**Table 2**). A relatively low fraction of sites on some arrays are polymorphic in NeuroGAP-Psychosis (e.g. only 422,156 of ~690,000 sites on GSA are polymorphic). We confirmed this expectation by calculating the mean proportion of SNPs at various frequencies on several GWAS arrays across 1000 Genomes populations (**Figure 3**).

**Figure 3 –.**
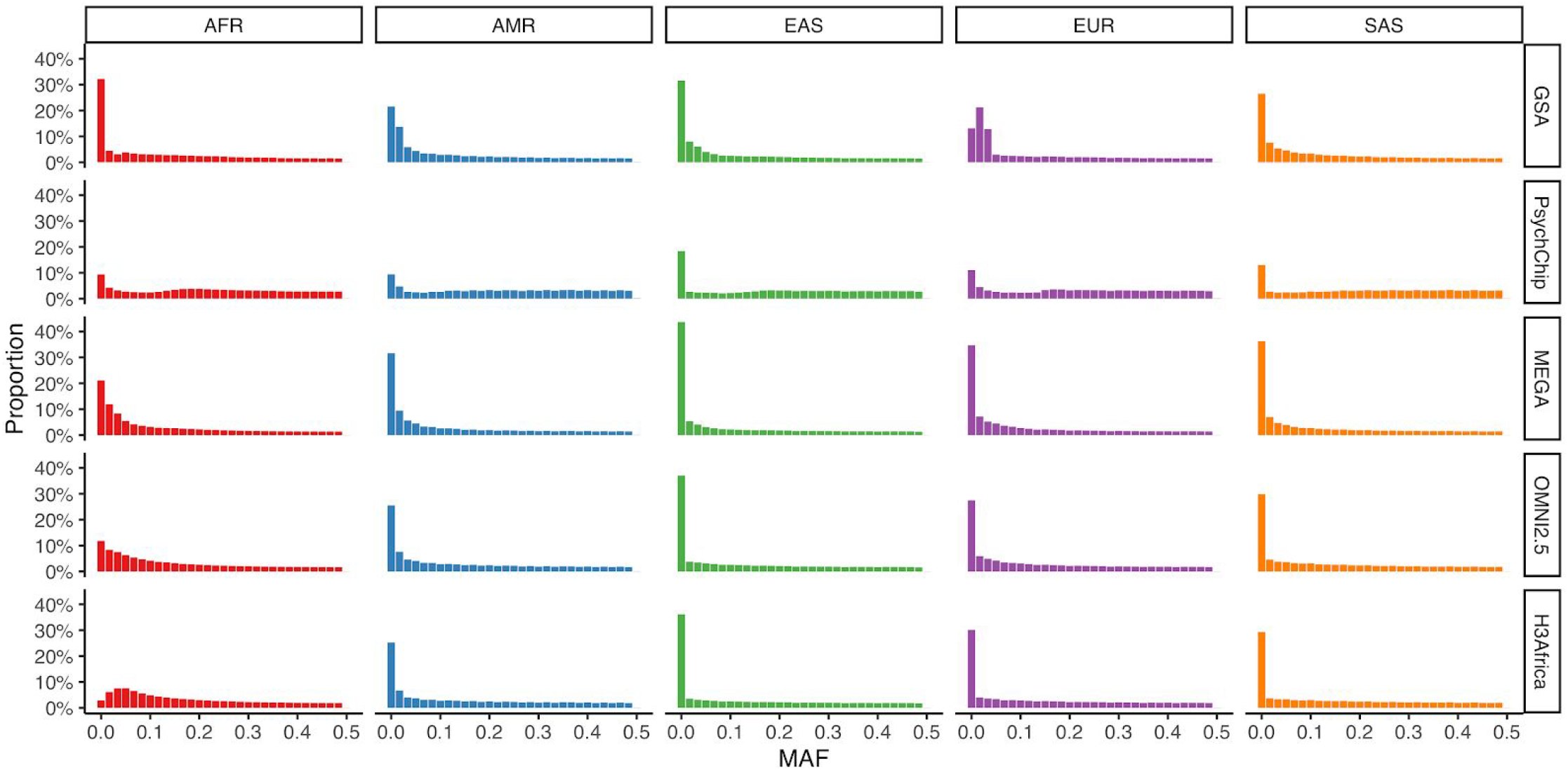
Minor allele frequency (MAF) across GWAS arrays and continental ancestries using 1000 Genomes data. AFR=Africans, AMR=admixed Americans (e.g. Hispanics/Latinos), EAS=East Asians, EUR=Europeans, and SAS=South Asians. These results indicate that the GSA captures variants that are especially common in Europeans relative to elsewhere.

**Table 2 –.** Sensitivity of various sequencing depths and GWAS arrays to detect singletons and common variants through several analytical steps. All numbers reported here are from processing using BEAGLE. Common variants here are defined as having > 5 copies (i.e. MAF>3%).

|  | # of SNPs |  |  | % singletons present in full set |  |  | % common variants in full set |  |  |
| --- | --- | --- | --- | --- | --- | --- | --- | --- | --- |
| Call set | Raw | Refined | Imputed | Raw | Refined | Imputed | Raw | Refined | Imputed |
| 0.5X | 9,236,562 | 7,452,675 | 18,414,145 | 0.04 | 0.01 | 0.33 | 0.55 | 0.40 | 0.62 |
| 1X | 13,036,891 | 10,389,726 | 18,974,677 | 0.09 | 0.03 | 0.33 | 0.74 | 0.52 | 0.62 |
| 2X | 15,716,019 | 13,387,436 | 19,887,495 | 0.2 | 0.08 | 0.33 | 0.88 | 0.59 | 0.62 |
| 4X | 20,958,987 | 16,458,866 | 21,083,626 | 0.45 | 0.17 | 0.33 | 0.95 | 0.61 | 0.62 |
| 6X | 23,352,341 | 17,633,642 | 21,402,104 | 0.62 | 0.23 | 0.33 | 0.97 | 0.61 | 0.62 |
| 10X | 24,955,954 | N/A | N/A | 0.8 | N/A | N/A | 0.98 | N/A | N/A |
| 20X | 25,136,680 | N/A | N/A | 0.93 | N/A | N/A | 0.99 | N/A | N/A |
| All reads | 26,093,644 | N/A | N/A | 1 | N/A | N/A | 1 | N/A | N/A |
| GSA | 422,156 | N/A | 18,272,172 | N/A | N/A | 0.33 | N/A | N/A | 0.62 |
| PsychChip | 350,678 | N/A | 18,190,171 | N/A | N/A | 0.33 | N/A | N/A | 0.62 |
| MEGA | 1,152,178 | N/A | 19,219,473 | N/A | N/A | 0.33 | N/A | N/A | 0.62 |
| H3Africa | 2,151,137 | N/A | 19,709,178 | N/A | N/A | 0.33 | N/A | N/A | 0.62 |
| Omni2.5 | 2,072,034 | N/A | 19,698,788 | N/A | N/A | 0.33 | N/A | N/A | 0.62 |

We also investigated the importance of the reference panel and the impact of missing population representation on sensitivity. For example, regardless of technology, we estimate that 33% of singletons in the “truth” dataset can be imputed (**Table 2**, i.e. 67% of singletons in the NeuroGAP-Psychosis data are absent or not tagged by the 1000 Genomes phase 3 data). This estimate is likely optimistic given that the low sample size in this study means that many variants reported here as singletons are likely somewhat common in the population. Additionally, 62% of common variants (AC > 5, MAF > 3%) in the “truth” dataset can be imputed, indicating that 38% of variants in the eastern and southern African populations in NeuroGAP-Psychosis are absent or untagged in the 1000 Genomes phase 3 data. While the number of variants imputed is inherently bounded by the reference data, the raw data indicates relatively high sensitivity to variants present in the “truth” data. For example, 45% of singletons in the full dataset can be detected with 4X data (**Table 2**). At the same depth, 95% of common variants are detected. As expected, we observe diminishing returns in numbers of variants imputed with increasing sequencing depth. More variants can be imputed with 2X sequencing using BEAGLE than with any of the GWAS arrays. Our sensitivity for detecting variants common in the truth data (74%) is higher with 1X sequencing than with imputed data from any array (62%, **Table 2**).

We next investigated variant call accuracy by calculating non-reference concordance across technologies. We also compared two imputation methodologies for use with low-coverage sequencing data--BEAGLE versus Gencove--as the latter was specifically designed for use with low-coverage data. Unlike BEAGLE, Gencove takes unmapped FASTQ files as an input to perform phasing and imputation, allowing consideration of genotype probabilities directly as described previously (Wasik et al., 2019). **Figure 4** shows non-reference concordance by allele frequency across sequencing versus array technologies, data processing steps through imputation with BEAGLE (“Refined” and “Imputed” panels in **Figure 4A**), in BEAGLE versus Gencove (**Figure 4B**), and in Gencove low-coverage versus GWAS array data imputed with BEAGLE (**Figure 4C**). **Figure 4A** includes different variants across panels including fewer but more accurate variants in the “Refined” panel, while the “Imputed” panel includes more than double the number of variants but with reduced accuracy (**Table 2**). When using BEAGLE for imputing both arrays and low-coverage data, these analyses indicate that the lower density arrays (GSA and PsychChip) perform similarly to 1X sequencing, medium density arrays (MEGA) perform almost as well as 2X data, and high density arrays (Omni2.5 array and H3Africa array specifically designed to capture African variation) perform between 2X and 4X sequencing (**Figure 4A**). We also compared the accuracy of two imputation methods, BEAGLE and Gencove, using the same set of imputed sites in the low-coverage sequencing data. We find that imputation performs better with Gencove for the lowest depths (0.5X, 1X, and 2X), while BEAGLE performs better for higher depths (4X and 6X, **Figure 4B, Supplementary Table 4**). When comparing low-coverage data imputed with Gencove versus GWAS array data imputed with BEAGLE, we see that 1X sequencing outperforms the low and medium density arrays (MEGA, GSA, and PsychChip), and that the high density arrays (H3Africa and Omni2.5) perform comparably to 2X sequencing (**Figure 4C, Supplementary Table 4**). Overall, these results show that GWAS arrays perform at best comparably to low-coverage sequencing.

**Figure 4 –.**
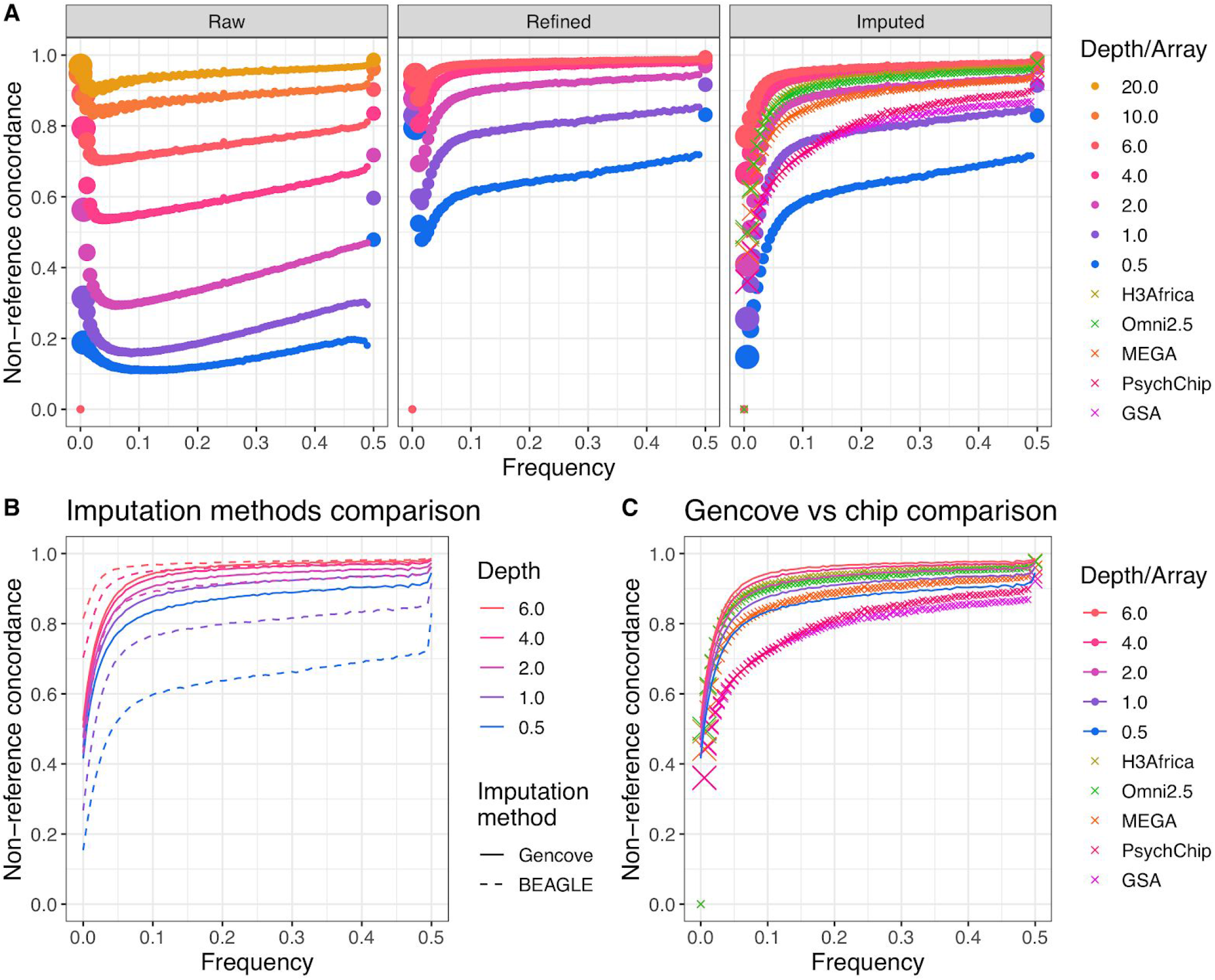
Non-reference concordance for SNPs as a function of sequencing depth or genotyping array, frequency, analysis stage, and imputation method. “Truth” dataset here is the full depth joint called sequencing dataset. All depths of sequencing data are shown for the raw data. Sequencing at 10X and 20X are excluded for all except the raw data because of minimal potential accuracy gains and to reduce computational costs. A) Non-reference concordance comparisons throughout steps of the BEAGLE analysis pipeline. Size of the points are proportional to the number of SNPs in each frequency bin. B) Non-reference concordance comparisons of BEAGLE versus Gencove for imputation of low coverage data. C) Non-reference concordance comparison of Gencove for imputation of low coverage data versus BEAGLE for imputation of GWAS arrays. Non-reference concordance values averaged across B-C) are shown in **Supplementary Table 4.**

We next investigated the impact of ancestral diversity on imputation accuracy from arrays versus sequencing depth. The populations in NeuroGAP-Psychosis span a broad range of geographical, ethnolinguistic, and ancestral diversity in eastern and southern Africa. Despite this considerable diversity with a range of genetic distances from populations represented in the 1000 Genomes reference haplotypes, there is remarkable qualitative consistency in data quality from various sequencing depths and GWAS arrays (**Figure 5**). We quantify subtle differences across populations (**Supplementary Table 5**). For example, imputation is least accurate among participants from Addis Ababa, Ethiopia. In contrast, imputation performs best in participants from Kilifi, Kenya, where some participants self-identify as Luhya. These differences in imputation accuracy across populations likely reflect genetic distances between the NeuroGAP-Psychosis participants and the 1000 Genomes phase 3 reference data, which includes for example a Luhya population from Kenya. These findings consistently indicate that 4X sequencing data outperforms all common commercial GWAS arrays for diverse African ancestry populations, including those specifically designed with African variation in mind such as the H3Africa array.

**Figure 5 –.**
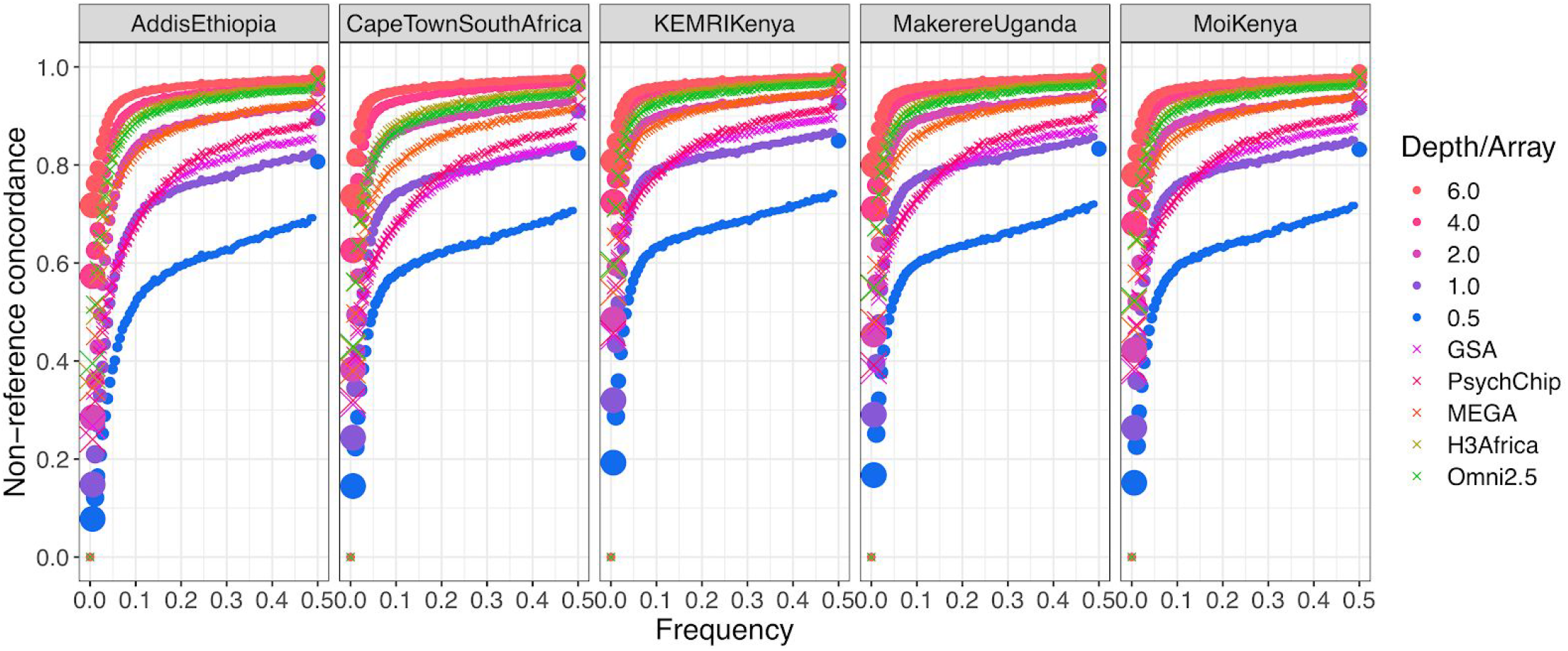
Non-reference concordance between imputed versus truth data across various populations and sites in Africa. Size of the points where applicable are proportional to the number of SNPs in each frequency bin. Quantitative comparisons across all variants and imputation methods are shown in **Supplementary Table 5**.

Depending on the tissue from which DNA is derived, another potential advantage of low-coverage sequencing over GWAS arrays is the ability to use off-target reads that do not map to *Homo sapiens* for further microbiome analysis. DNA from this study is derived from saliva, enabling oral microbiome quantifications. We used taxonomic profiling quantifications from the software Kraken, which were produced from 6X data input to Gencove. For each individual, we quantified relative abundances from read counts. We show the phylum-level relative abundances as a proof-of-concept (**Supplementary Figure 3**).

Lastly, we list realistic pricing for low-coverage sequencing versus GWAS arrays based on current publicly available reagent costs from Illumina (**Table 3**). While these do not include fixed sample and library preparation costs, we assume that these are comparable across GWAS arrays and sequencing approaches. We note that all costs can vary considerably depending on consortium pricing, sequencing facility, volume, etc. While sequencing costs list volume discounts (e.g. up to 39% discount for high volume flow cell purchasing), GWAS arrays do not; to compare these technologies as fairly as possible, we therefore list the non-discounted price but note that costs could be lower (**Supplementary Table 6**). Based on these prices, we show that the high-density arrays are similar in price to 4-6X sequencing. The lowest depths of sequencing evaluated here, 0.5-1X, are cheaper than the PsychChip and GSA.

**Table 3 –.** Costs of reagents for sequencing and genotyping options. We aggregated the prices for reagents from Illumina’s website as of April 10, 2020. These prices notably do not include sample and library preparation costs, which we assume to be comparable between GWAS arrays and sequencing approaches. The H3Africa array is not commercially listed on Illumina’s site and is thus not included here. Sequencing reagent costs use Illumina’s list price for the NovaSeq 6000 S4 Reagent Kit. Each flow cell has a maximum output of 3,000 Gb. Prices listed below assume single flow cell purchasing, which is listed at $31,700. Prices adjusting for bulk flow cell purchasing from Illumina are shown in **Supplementary Table 6**. Sequencing costs assume 125 Gb to achieve a target depth of 30X whole genome sequencing coverage.

| Depth/Array | Reagent cost per sample |
| --- | --- |
| 30X | \$1,320.83 |
| 20X | \$880.55 |
| 6X | \$264.17 |
| Omni2.5 | \$184.43 |
| 4X | \$176.11 |
| MEGA Global | \$119.00 |
| 2X | \$88.06 |
| PsychChip | \$71.38 |
| GSA | \$49.00 |
| 1X | \$44.03 |
| 0.5X | \$22.01 |

Another pricing consideration regarding different depths of sequencing or GWAS arrays is the computational complexity. Genotype refinement is only necessary for low-coverage sequencing and is a more computationally complex step than imputation. Imputation is also slightly more costly with low-coverage sequencing than with GWAS arrays because more variants are called from the beginning, increasing genomic coverage. However, we find that the computational costs of genotype refinement and the slightly increased computational complexity of imputation from more variants called at the outset are negligible compared to data generation costs. For low-coverage sequencing, reagent costs alone are ≥100 times higher than the sum of refinement and imputation depending on depth of coverage (ratio increasing with higher depths), and GWAS array costs are >2800 times higher than imputation (ratio increasing with higher array density, **Supplementary Table 7**).

## Discussion

In this study, we have compared the relative merits and costs of several genetic data generation and processing strategies in a diverse cohort of eastern and southern Africans. We conclude that 4X sequencing outperforms all GWAS arrays evaluated, including dense arrays. This outcome is in spite of the fact that the dense H3Africa array was designed to capture African variation and thus tags the most variation in the NeuroGAP-Psychosis data of all GWAS arrays analyzed here. 4X sequencing is comparable in price to high-density arrays that assay millions of SNPs and indels across the allele frequency spectrum. Among more affordable options, we find that 0.5-1X sequencing costs less than and performs similarly to or better than commonly used lower density arrays such as the Illumina GSA. Additionally, we note that the GSA is composed of variants most common in European populations and is thus not the most appropriate technology for studies of participants with primarily non-European ancestry.

Low-coverage sequencing has several distinct advantages compared to GWAS arrays. It facilitates the identification of novel variation across the allele frequency spectrum. This is especially valuable in non-European populations where haplotype reference panels used for imputation are currently lacking, as it provides the opportunity to construct them. In these NeuroGAP-Psychosis data, we find that 38% of common variants could not be imputed from the 1000 Genomes phase 3 data, likely due to a dearth of eastern and southern African diversity represented in this reference panel. Among rare variants, we find that 4X sequencing detects nearly half of all singletons, an especially appealing attribute for studies of diseases that have undergone negative selection. We plan to build on previous work in European ancestry populations by the Haplotype Reference Consortium, which aggregated low-coverage sequencing data into an imputation panel (McCarthy et al., 2016). Specifically, the high coverage genomes sequenced here along with additional low-coverage whole genomes will be integrated into a more diverse reference panel to improve phasing and imputation. In contrast to GWAS arrays, sequencing also offers the opportunity to use off-target reads to measure microbiome variation, as demonstrated here with saliva-derived DNA. Additional post-GWAS methodological advances are possible from low-coverage sequencing as well; for example, pooling reads from controls separately from cases that contribute to GWAS peaks can enhance variant discovery, an important step for fine-mapping.

While GWAS arrays are most commonly used in current large-scale genetic studies, low-coverage sequencing has several distinct advantages in populations underrepresented in genomics, especially in African populations where overall genetic variation is higher, linkage disequilibrium is shorter, and haplotype reference data are lacking. Despite African populations having the most genetic variation globally, with as much variation among individuals from different regions of Africa as between some continents, African ancestry genomes are vastly underrepresented. Further, the vast majority of African ancestry participants in genetic studies are African Americans or Afro-Caribbeans (72-93% in the GWAS catalog and ≥90% in gnomAD) with primarily West African ancestors (Martin et al., 2018). In addition to informing the most appropriate and cost-effective data generation strategies, this study also adds to a relatively small number of high coverage whole genomes sequenced from Africa.

## Methods

### Human subjects

Ethical and safety considerations are being taken across multiple levels, as described in greater detail previously (Stevenson et al., 2019). Since the subjects the study aims to recruit are deemed vulnerable populations, additional measures are taken to protect them. Potential participants are excluded if they are presenting with severe, intrusive levels of psychiatric symptoms at the time of consent. Additionally, researcher assistants use the University of California, San Diego Brief Assessment of Capacity to Consent (UBACC) system (Campbell et al., 2017; Jeste et al., 2007) during the consent process to make sure participants understand the study, what is required of them, and that they can withdraw at any point. Participants who pass the UBACC and who want to continue are required to provide written informed consent or a fingerprint in lieu of a signature. No protected health information (PHI) or Health Insurance Portability and Accountability Act (HIPPA) identifiers are collected as part of the phenotypic or genetic dataset.

Ethical clearances to conduct this study have been obtained from all participating sites, including:

- Ethiopia: Addis Ababa University College of Health Sciences (#014/17/Psy) and the Ministry of Science and Technology National Research Ethics Review Committee (#3.10/14/2018).
- Kenya: Moi University College of Health Sciences/Moi Teaching and Referral Hospital Institutional Research and Ethics Committee (IREC) (#IREC/2016/145, approval number: IREC 1727), Kenya National Council of Science and Technology (#NACOSTI/P/17/56302/19576) KEMRI Centre Scientific Committee (CSC# KEMRI/CGMRC/CSC/070/2016), KEMRI Scientific and Ethics Review Unit (SERU# KEMRI/SERU/CGMR-C/070/3575)
- South Africa: The University of Cape Town Human Research Ethics Committee (#466/2016)
- Uganda: The Makerere University School of Medicine Research and Ethics Committee (SOMREC #REC REF 2016-057) and the Uganda National Council for Science and Technology (UNCST #HS14ES)
- USA: The Harvard T.H. Chan School of Public Health (#IRB17-0822)

### Variant calling

We used the GATK best practices pipeline described for variant calling using code available here: https://github.com/gatk-workflows/gatk4-germline-snps-indels. For most steps, we used GATK4 with Cromwell (https://cromwell.readthedocs.io/en/stable/) to submit jobs in parallel across the genome where possible using the Google Cloud Platform.

### Depth of coverage

Depth statistics from high coverage whole genomes were computed by the Broad Institute’s Data Science Platform team. This calculation excluded low quality, unmapped, unpaired, and duplicate reads in depth of coverage calculations.

### Downsampling sequencing reads

We downsampled reads using the GATK DownsampleSam module, which retains a deterministically random subset of reads and their mate pairs. We calculated the probability used for downsampling based on depth of coverage as described above (i.e. not simply based on the total number of reads sequenced relative to the number of bases in the human genome because for example some reads from saliva-derived DNA may not be human).

### Concordance

We computed non-reference concordance among homozygous reference, heterozygous, and homozygous non-reference calls, excluding no call and missing sites from counts, according to the following table:

|  | No call | ./. | 0/0 | 0/1 | 1/1 |
| --- | --- | --- | --- | --- | --- |
| No call | a | b | c | d | e |
| ./. | f | g | h | i | j |
| 0/0 | k | l | m | n | o |
| 0/1 | p | q | r | s | t |
| 1/1 | u | v | w | x | y |

Concordance = s + y / (n + o + r + s + t + w + x + y)

Homozygous reference concordant calls were excluded to avoid high concordance among rarer variants by simply imputing the most common allele.

### Haplotype reference

We downloaded phased 1000 Genomes haplotype reference data containing SNPs aligned to GRCh38 from the following location: ftp://ftp.1000genomes.ebi.ac.uk/vol1/ftp/data_collections/1000_genomes_proiect/release/20181203_biallelic_S_NV/. We used these phased haplotypes for genotype refinements, phasing, and imputation.

### Genotype refinement, phasing, and imputation

We used BEAGLE 4.1 for genotype refinement of variant calls in downsampled sequencing data prior to phasing and imputation. We then used BEAGLE 5.1 for phasing and imputation both for variant call corresponding to low-coverage sequencing data and GWAS array data.

### Gencove imputation

We generated FASTQ files from analysis-ready BAM files using the bedtools bamtofastq. We then uploaded these FASTQ files to the Gencove server, ran imputation and related analyses, then downloaded imputation results.

## Acknowledgments

We thank Juha Karjalainen for his help setting up and troubleshooting a Cromwell server for running all workflows. We thank Laura Gauthier for explanations of components of the Broad Institute Data Science Platform pipelines.

## Authors’ contributions

A.R.M. conceived of the study, conducted analyses, and wrote the manuscript. E.G.A. provided feedback on the manuscript. M.J.D., B.M.N, and E.G.A. provided analytical input. J.K.P. provided analytical tools. S.B.C, A.S., R.E.S., and C.N. managed data intake and coordinated project management across sites. T.B., T.D., S. D., and S.F. helped with sequencing logistics and cost analyses. C.v.d.M., D.S. helped with cost analyses. T. A., M.A., F.A., S.G., W.E.I., R.J., G.K., E.K., J.K., L.M., H.M., R.M.M., C.N., W.S., A.S., R.E.S., and R.R. were responsible for data acquisition. D.A., L.A., M.J.D., B.G., S.K., K.C.K, B.M.N., C.R.J.C.N., D.J.S., S.T., and Z.Z. oversaw the study.

## Web resources

Code used to process and analyze data is available here: https://github.com/armartin/neurogap_downsampling.

