## Supplementary Figures 1-3, Tables 1-7 for "Low-coverage sequencing cost-effectively detects known and novel variation in underrepresented populations"

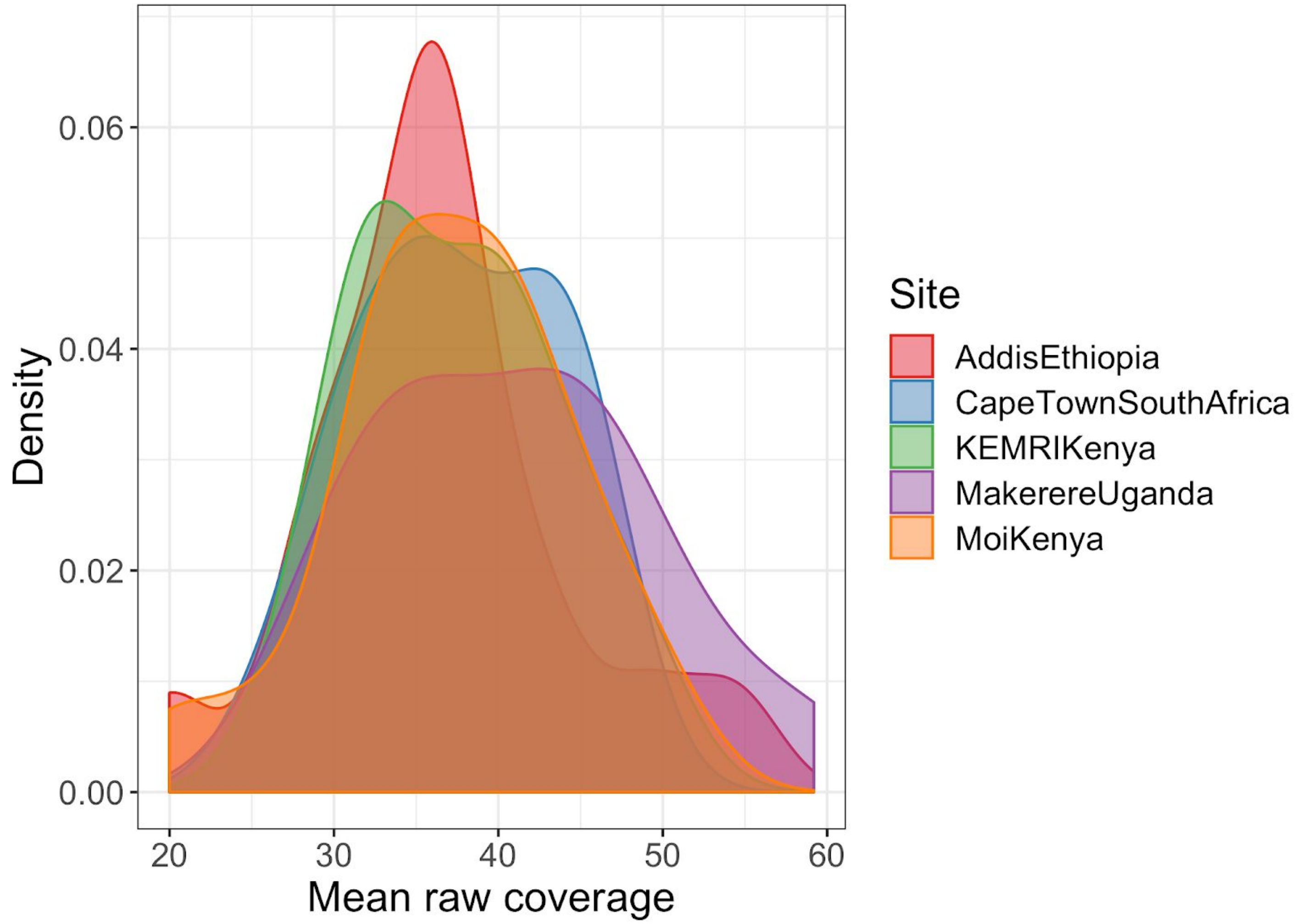


**Supplementary Figure 1 - Mean coverage across 91 NeuroGAP whole genomes.**

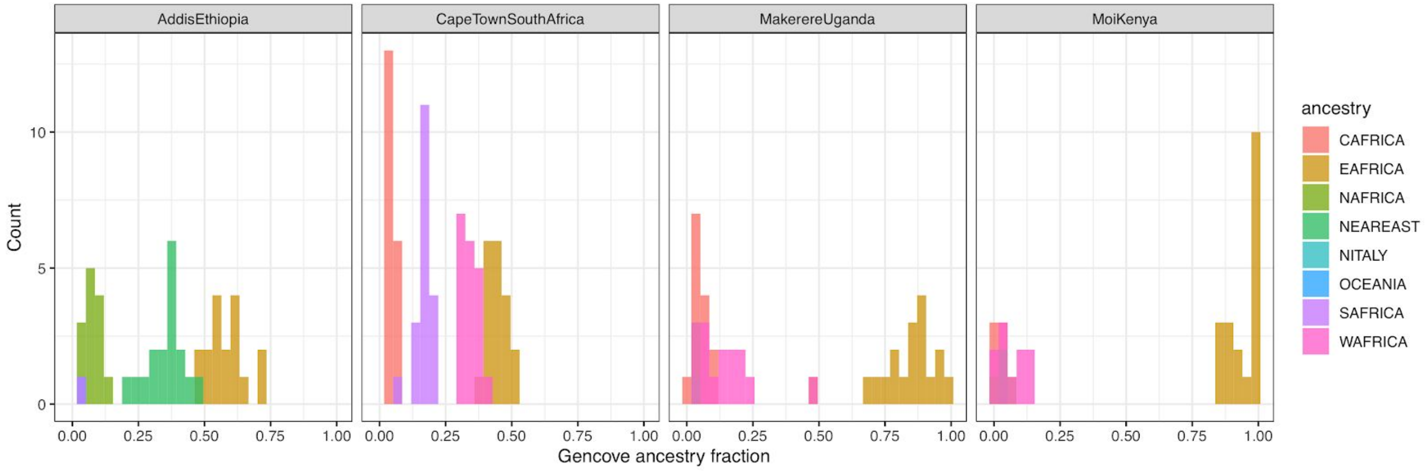


**Supplementary Figure 2 - Ancestry report generated by Gencove.**

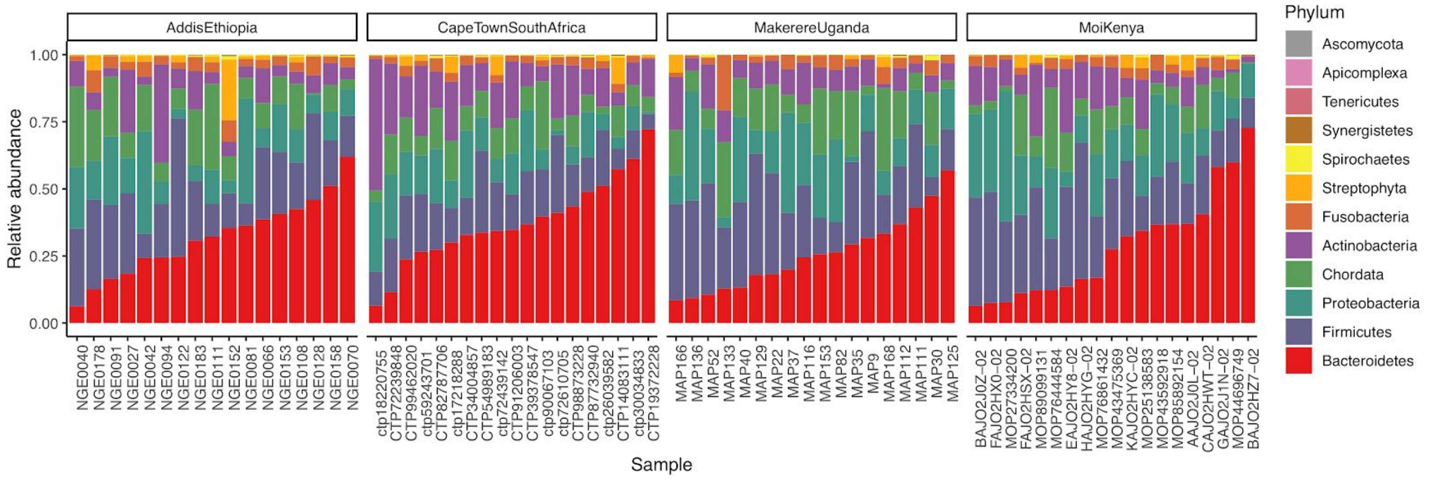


**Supplementary Figure 3 - Phylum-level microbiome variation.** Results were generated using Kraken 1.0 with unmapped reads from the 6X coverage downsampled data. All phyla with a non-zero relative abundance across individuals are shown, in order of mean abundance (i.e. bacteroidetes on bottom have the highest mean relative abundance), and individuals are ordered within site based on their relative abundance of the most frequent phylum.

**Supplementary Table 1 - Counts of primary self-reported ethnicities by project site**

| **NeuroGAP site** | **Primary ethnicity** | **Count** |
|---|---|---|
| AddisEthiopia | Amhara | 9 |
| AddisEthiopia | Oromo | 6 |
| AddisEthiopia | Sebat Bet Gurage | 1 |
| AddisEthiopia | Sidama | 1 |
| AddisEthiopia | Silt'e | 1 |
| AddisEthiopia | Sodo Gurage | 1 |
| CapeTownSouthAfrica | Xhosa | 17 |
| CapeTownSouthAfrica | Other (please specify) | 1 |
| CapeTownSouthAfrica | Zulu | 1 |
| KEMRIKenya | Mijikenda | 10 |
| KEMRIKenya | Kamba | 2 |
| KEMRIKenya | Luhya | 2 |
| KEMRIKenya | Chonyi | 1 |
| KEMRIKenya | Giriama | 1 |
| KEMRIKenya | Meru | 1 |
| KEMRIKenya | Other (please specify) | 1 |
| MakerereUganda | Baganda | 3 |
| MakerereUganda | Lugbara | 3 |
| MakerereUganda | Banyankore | 2 |
| MakerereUganda | Basoga | 2 |
| MakerereUganda | Iteso | 2 |

| | | |
|---|---|---|
| MakerereUganda | Bafumbira | 1 |
| MakerereUganda | Bakonzo | 1 |
| MakerereUganda | Banyoro | 1 |
| MakerereUganda | Karimojong | 1 |
| MakerereUganda | Madi | 1 |
| MakerereUganda | Sabiny | 1 |
| MoiKenya | Kalenjin | 9 |
| MoiKenya | Kikuyu | 4 |
| MoiKenya | Luhya | 4 |
| MoiKenya | Luo | 2 |

**Supplementary Table 2 - Sensitivity and quality control metrics from downsampling experiment using raw variant call metrics.** The metrics at the top of the table (TOTAL_SNPS through NUM_SINGLETONS) were produced by the Picard software. Values in the lower rows were produced by custom scripts (**Web Resources**). Common variants here are defined as having > 5 copies (i.e. MAF>3%).

| **Depth** | **0.5X** | **1X** | **2X** | **4X** | **6X** | **10X** | **20X** | **All reads** |
|---|---|---|---|---|---|---|---|---|
| **TOTAL_SNPS** | 9,236,562 | 13,036,891 | 15,716,019 | 20,958,987 | 23,352,341 | 24,955,954 | 25,136,680 | 26,093,644 |
| **PCT_DBSNP** | 0.81 | 0.79 | 0.83 | 0.77 | 0.74 | 0.72 | 0.73 | 0.71 |
| **DBSNP_TITV** | 2.11 | 2.13 | 2.15 | 2.16 | 2.17 | 2.18 | 2.18 | 2.18 |
| **NOVEL_TITV** | 1.6 | 1.6 | 1.84 | 1.92 | 1.95 | 1.98 | 1.93 | 1.9 |
| **TOTAL_INDELS** | 1,330,023 | 1,813,310 | 2,382,243 | 2,962,429 | 3,311,102 | 3,269,766 | 3,033,225 | 3,034,130 |
| **PCT_DBSNP_INDELS** | 0.77 | 0.68 | 0.58 | 0.49 | 0.45 | 0.46 | 0.5 | 0.5 |
| **DBSNP_INS_DEL_RATIO** | 0.81 | 0.76 | 0.7 | 0.67 | 0.66 | 0.65 | 0.66 | 0.66 |
| **NOVEL_INS_DEL_RATIO** | 0.51 | 0.48 | 0.41 | 0.37 | 0.39 | 0.49 | 0.63 | 0.66 |
| **TOTAL_MULTIALLELIC_SNPS** | 51,827 | 114,941 | 193,097 | 324,576 | 395,427 | 458,749 | 471,974 | 406,266 |
| **NUM_IN_DB_SNP_MULTIALLELIC** | 44,922 | 94,856 | 152,005 | 237,526 | 277,126 | 307,149 | 305,615 | 264,302 |
| **TOTAL_COMPLEX_INDELS** | 195,879 | 414,268 | 625,125 | 828,820 | 996,225 | 1,117,219 | 1,211,503 | 1,238,754 |
| **NUM_IN_DB_SNP_COMPLEX_INDELS** | 182,848 | 375,943 | 544,172 | 684,130 | 778,092 | 833,033 | 867,798 | 876,455 |
| **SNP_REFERENCE_BIAS** | 0.38 | 0.38 | 0.41 | 0.45 | 0.47 | 0.5 | 0.5 | 0.51 |
| **NUM_SINGLETONS** | 1,161,967 | 2,215,593 | 3,777,977 | 7,040,205 | 8,697,345 | 9,579,361 | 9,264,341 | 9,505,281 |
| **n_hom_ref (mean)** | 3,171,751 | 7,898,931 | 14,346,304 | 23,134,834 | 26,670,673 | 28,641,510 | 28,468,205 | 31,926,975 |
| **n_het (mean)** | 45,177 | 170,224 | 621,024 | 1,795,694 | 2,720,851 | 3,630,054 | 4,109,947 | 4,148,694 |
| **n_hom_alt (mean)** | 322,066 | 736,349 | 1,414,176 | 1,984,122 | 2,056,318 | 2,013,028 | 1,986,839 | 1,924,021 |
| **Fraction singletons present in full set** | 0.04 | 0.09 | 0.2 | 0.45 | 0.62 | 0.8 | 0.93 | 1 |

| | | | | | | | | |
|---|---|---|---|---|---|---|---|---|
| **Fraction of 2-5 copy sites in full set** | 0.09 | 0.21 | 0.42 | 0.7 | 0.81 | 0.88 | 0.94 | 1 |
| **Fraction common variants in full set** | 0.55 | 0.74 | 0.88 | 0.95 | 0.97 | 0.98 | 0.99 | 1 |
| **Genome-wide concordance** | 0.22 | 0.42 | 0.66 | 0.86 | 0.93 | 0.97 | 0.98 | 1 |

**Supplementary Table 3 - Raw SNP and indel non-reference variant concordance from low-coverage genomes with full coverage genomes prior to genotype refinement or imputation.** Concordance is averaged across variants of all allele frequencies.

| **Depth** | **SNPs** | **Indels** |
|---|---|---|
| 0.5X | 0.12 | 0.10 |
| 1X | 0.17 | 0.12 |
| 2X | 0.30 | 0.19 |
| 4X | 0.54 | 0.35 |
| 6X | 0.70 | 0.49 |
| 10X | 0.84 | 0.65 |
| 20X | 0.91 | 0.83 |

**Supplementary Table 4 - Non-reference concordance across methods and technologies.** Values reported are across all SNPs shown in **Figure 4**.

| **Depth/array** | **Method** | **Overall non-reference concordance** |
|---|---|---|
| 6X | BEAGLE | 0.975 |
| 4X | BEAGLE | 0.959 |
| 6X | Gencove | 0.949 |
| 4X | Gencove | 0.94 |
| H3Africa | BEAGLE | 0.932 |
| Omni2.5 | BEAGLE | 0.926 |
| 2X | Gencove | 0.924 |
| 2X | BEAGLE | 0.91 |
| 1X | Gencove | 0.904 |
| MEGA | BEAGLE | 0.892 |

| | | |
|---|---|---|
| 0.5X | Gencove | 0.875 |
| PsychChip | BEAGLE | 0.829 |
| GSA | BEAGLE | 0.816 |
| 1X | BEAGLE | 0.815 |
| 0.5X | BEAGLE | 0.681 |

**Supplementary Table 5 - Average non-reference concordance across technologies and allele frequencies for each population.** Values for each site show non-reference concordance. The same imputation reference panel, 1000 Genomes phase 3 data, was used for all analyses, including as input to both BEAGLE and Gencove.

| Depth/array | Method | AddisEthiopia | CapeTownSouthAfrica | KEMRIKenya | MakerereUganda | MoiKenya |
|---|---|---|---|---|---|---|
| 6X | BEAGLE | 0.961 | 0.964 | 0.971 | 0.97 | 0.968 |
| 4X | BEAGLE | 0.939 | 0.946 | 0.958 | 0.956 | 0.953 |
| H3Africa | BEAGLE | 0.922 | 0.91 | 0.949 | 0.941 | 0.94 |
| Omni2.5 | BEAGLE | 0.918 | 0.902 | 0.944 | 0.935 | 0.934 |
| 6X | Gencove | 0.908 | 0.909 | N/A | 0.929 | 0.927 |
| 4X | Gencove | 0.899 | 0.9 | N/A | 0.923 | 0.92 |
| 2X | Gencove | 0.883 | 0.882 | N/A | 0.911 | 0.908 |
| 2X | BEAGLE | 0.877 | 0.892 | 0.919 | 0.913 | 0.907 |
| MEGA | BEAGLE | 0.88 | 0.861 | 0.918 | 0.901 | 0.902 |
| 1X | Gencove | 0.862 | 0.862 | N/A | 0.894 | 0.891 |
| 0.5X | Gencove | 0.831 | 0.833 | N/A | 0.869 | 0.865 |
| PsychChip | BEAGLE | 0.808 | 0.798 | 0.864 | 0.836 | 0.839 |
| GSA | BEAGLE | 0.796 | 0.782 | 0.854 | 0.822 | 0.827 |
| 1X | BEAGLE | 0.771 | 0.795 | 0.835 | 0.82 | 0.813 |
| 0.5X | BEAGLE | 0.639 | 0.663 | 0.709 | 0.683 | 0.68 |

**Supplementary Table 6 - Costs of reagents for sequencing and genotyping options including sequencing volume discounts.** We aggregated list prices of reagents from Illumina’s website as of April 10, 2020. These prices notably do not include sample and library preparation costs, which we assume to be comparable between GWAS arrays and sequencing approaches. The H3Africa array is not commercially listed on Illumina’s site and is thus not included here. Sequencing reagent costs assume Illumina’s list price of the NovaSeq 6000 S4 Reagent Kit. Bulk pricing listed at $240,000 for 10 flow cells ($24,000/flow cell), $456,000 for 20 flow cells ($22,800/flow cell), and $768,000 for 40 flow cells ($19,200/flow cell) reduces costs at large scales. Rows are sorted based on the largest bulk purchasing cost.

| Depth/Array | List cost | Bulk purchasing (10 flow cells for $240,000) | Bulk purchasing (20 flow cells for $456,000) | Bulk purchasing (40 flow cells for $768,000) |
|---|---|---|---|---|
| 30X | 1,320.83 | $1,000.00 | $950.00 | $800.00 |
| 20X | $880.55 | $666.67 | $633.33 | $533.33 |
| Omni2.5 | $184.43 | $184.43 | $184.43 | $184.43 |
| 6X | $264.17 | $200.00 | $190.00 | $160.00 |
| MEGA Global | $119.00 | $119.00 | $119.00 | $119.00 |
| 4X | $176.11 | $133.33 | $126.67 | $106.67 |
| PsychChip | $71.38 | $71.38 | $71.38 | $71.38 |
| 2X | $88.06 | $66.67 | $63.33 | $53.33 |
| GSA | $49.00 | $49.00 | $49.00 | $49.00 |
| 1X | $44.03 | $33.33 | $31.67 | $26.67 |
| 0.5X | $22.01 | $16.67 | $15.83 | $13.33 |
| H3Africa | Unknown | Unknown | Unknown | Unknown |

**Supplementary Table 7 - Compute times for genotype refinement and imputation using BEAGLE.** We assume a computational cost of $0.02 / CPU hour run on custom machines with 11 Gb of RAM as these were run across ~1000 shards on Google Cloud preemptible nodes. Costs were divided across 93 samples, 2 of which were dropped from analysis due to contamination. Some values are missing because job failures required multiple iterations of resubmissions.

| **Depth/Array** | **Step** | **Total run time (s)** | **Cost per sample** |
|---|---|---|---|
| 0.5X | Refinement | 3218175 | $0.19 |
| 1X | Refinement | 5443643 | $0.33 |
| 2X | Refinement | 8962256 | $0.54 |
| 4X | Refinement | 14103035 | $0.84 |
| 0.5X | Imputation | 576078 | $0.03 |
| 1X | Imputation | 536017 | $0.03 |
| 2X | Imputation | 581023 | $0.03 |
| Omni2.5 | Imputation | 381759 | $0.02 |
| H3Africa | Imputation | 362223 | $0.02 |
| MEGA | Imputation | 326611 | $0.02 |
| PsychChip | Imputation | 292045 | $0.02 |
| GSA | Imputation | 287468 | $0.02 |
